# Multi-Template Matching: a versatile tool for object-localization in microscopy images

**DOI:** 10.1101/619338

**Authors:** Laurent S. V. Thomas, Jochen Gehrig

## Abstract

We implemented multiple template matching as both a Fiji plugin and a KNIME workflow, providing an easy-to-use method for the automatic localization of objects of interest in images. We demonstrate its application for the localization of entire or partial biological objects. The Fiji plugin can be installed by activating the Multi-Template-Matching and IJ-OpenCV update sites. The KNIME workflow can be downloaded from <u>nodepit space</u> or the associated GitHub repository. Python source codes and documentations are available on the following GitHub repositories: <u>LauLauThom/MultiTemplateMatching</u> and <u>LauLauThom/MultipleTemplateMatching-KNIME</u>.

## Background

In microscopy images, the objects of interest usually represent only a fraction of the field of view and are randomly positioned. While classical image processing methods like thresholding followed by binary operations and identification of connected components perform well for the localization of fluorescently-labelled objects, this approach often requires substantial engineering and results in highly application-specific solutions. In many cases, such as whole organism imaging, such methods might not even be applicable when it comes to the identification of a particular organ or tissue that is poorly contrasted from the rest of the specimen or that is non-specifically labelled. Machine learning methods offer powerful detection capacities to overcome such challenges (Gallego *et al.*, 2018; Waithe *et al.*, 2019; Falk *et al.*, 2019); however, their implementation and the training of the machine is usually complex and requires large amounts of data, thus rendering it often inaccessible to most microscopy users. In contrast, template-based approaches allow the computation of the most probable positions of a single template image within a larger image with negligible manual annotation and at minimal computational cost. Here, we demonstrate the implementation of template-matching in Fiji (Schindelin *et al.*, 2012) and KNIME (Berthold *et al.*, 2009) in a user-friendly manner and its adaptation for the detection of objects in images, based on one or several user-provided templates.

## Implementation

We implemented template matching (Brunelli, 2009) as both a Fiji plugin and a KNIME workflow compatible with images of any bit depth. To predict the position of a template within a target image, the algorithm first computes a correlation map using the function *matchTemplate* from OpenCV, within Fiji (Domínguez *et al.*, 2017) or KNIME (Fig.1B, 1E and Suppl. Fig.1B). If information about the approximate position of the object is known a priori, the computation can be limited to a rectangular search region, which significantly speeds up the execution (Fig.1A,B; Suppl. Fig.6 and Suppl. Movie 1). To maximize the robustness of object-detection, our implementation allows to provide a set of templates to be searched, or the initial templates can be transformed by flipping and/or rotation. The computation of the correlation map is followed by maxima detection and Non-Maxima Suppression (NMS) (Alexe *et al.*, 2012; Felzenszwalb *et al.*, 2010) to prevent overlapping detections when several objects are expected (Suppl. Fig.1D,E). The detailed procedure for maxima detection and NMS can be found in Supplementary Material and a flowchart of the implementation is provided in Suppl. Fig.2. The parameters for the detection (template rotation/flipping, expected number of objects, score type and if N>1 threshold on the score and maximal overlap between predicted locations) are specified via a graphical user interface in both Fiji and KNIME (Suppl. Fig.3A, 4B). The detected regions are returned as rectangular regions of interest (ROI) in Fiji and as part of a mask image in KNIME along with a result table listing the score and the coordinates for each detection (Suppl. Fig.3B, 4C). The tools are intuitively accessible and demand no programming experience. Importantly, the Fiji plugin is macro-recordable, and can thus be readily integrated in custom image processing workflows (see Supplementary Macro for 2-step template matching). The resulting pipelines can be installed on any system running Fiji/KNIME.

**Figure 1:**
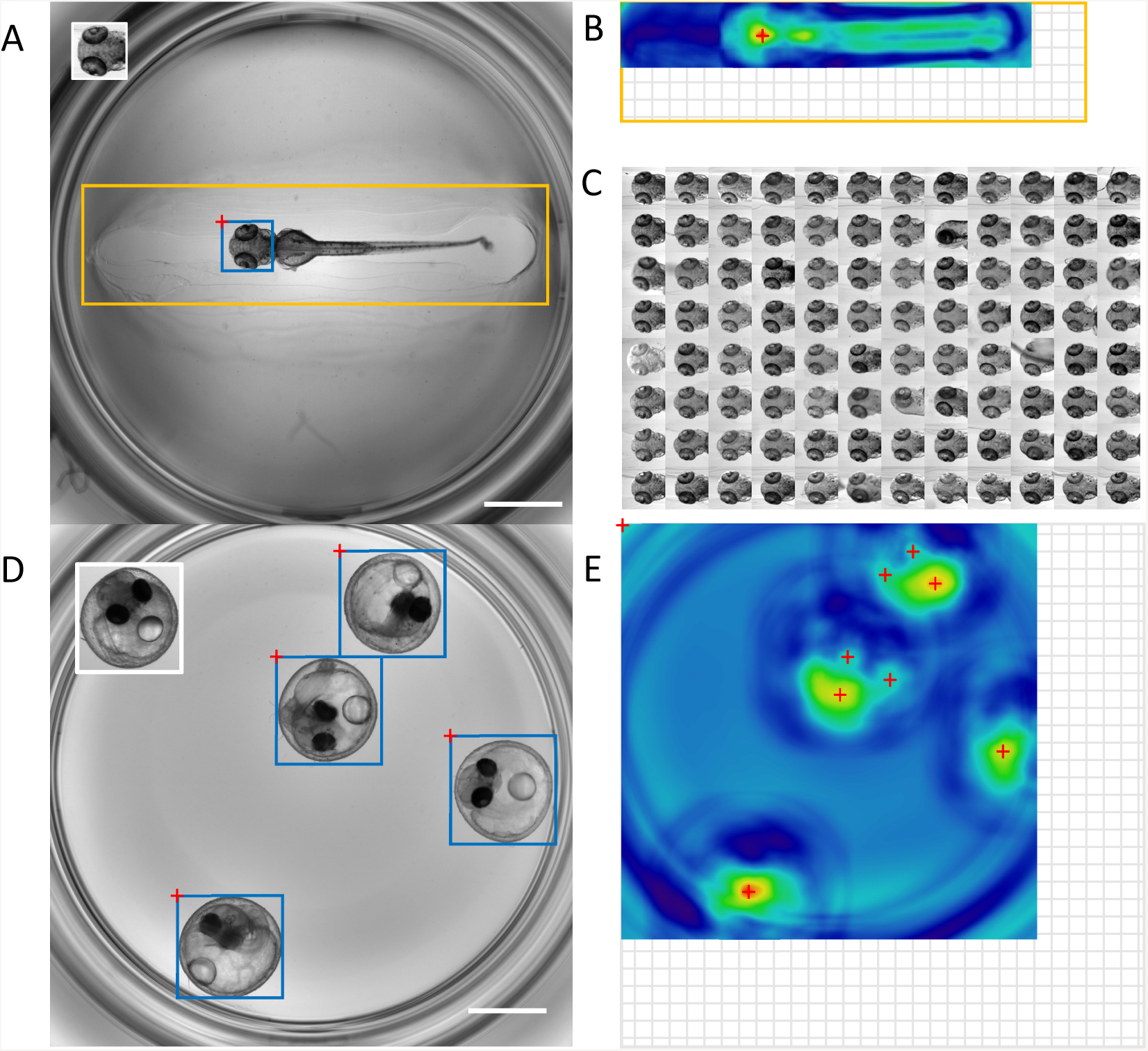
Single and multiple object detection by template matching. Template matching for head region detection in oriented zebrafish larvae with template as inset (188×194 pixels), search region in orange and predicted location in blue. (B) Correlation map: the red cross indicates the global maximum, which gives the position of the bounding box in A. (C) Montage of detected head regions for a 96 well plate. (D) Template matching for the detection of randomly oriented and positioned medaka embryos with inset showing initial template used (400×414 pixels), predicted object locations in blue and associated local maxima in red. In this case, the search is performed with the initial template as well as flipped (vertical and horizontal) and rotated versions of the initial and flipped templates (90,180,270°). (E) One of the correlation maps derived from D. The locations of possible local maxima before NMS is shown (red crosses). Image dimensions for A and D: 2048×2048 pixels with scale bar of 1 mm. Grid areas in B and E indicate the smaller size of the correlation map compared to the image in which the search is performed, as explained in Supplementary material.

## Results

We successfully tested our pipelines for the detection of organs like head and eyes in oriented zebrafish larvae, or for the localization of multiple randomly oriented and positioned medaka embryos (Gierten *et al.*, 2019) (Fig.1, Suppl. Fig.1, 5-7 and Suppl. Movie 1,2). The implemented method performs robustly, even with changes of specimen morphology as it often occurs e.g. in chemical screening scenarios, or with variations of illumination (Fig.1C, Suppl.Fig.7D and Suppl.Data). Our implementation relies on the OpenCV library which provides a rapid computation of the correlation map (about 0.45s/image in Fiji with a 188×194 template and 2048×2048 image on a laptop with an intel i7-7500U CPU). This computation can be even faster if a search region is provided (0.06 s/image with a 1820×452 search area as in Fig.1A,B and Suppl.Fig.5). When several templates are provided (e.g. additional geometrical transformations of the template or perspectives of the same object) the computations are repeated for each template which increases the computation time proportionally but typically increases the accuracy of the localization (Suppl. Fig.7). The implementation in KNIME is slower, because of the shared processing between the workflow and the local python installation but has improved modularity. Previous implementations of template matching (Peravali *et al.*, 2011; Tseng *et al.*, 2011; Domínguez *et al.*, 2017) do not allow to use different templates for object detections, neither do they provide a way to prevent multiple detections of the same object, which motivated the implementation of a NMS. Finally, the demonstrated template matching pipeline could also facilitate feedback microcopy applications by interfacing it with the control software of automated microscopes, thus enabling the automated acquisition of ROI for tracking or automated zooming-in on target structures without manual intervention (Peravali *et al.*, 2011; Pandey *et al.*, 2019).

## Conclusion

We demonstrate an implementation of template matching in Fiji and KNIME allowing the localization of one or several objects of interest based on template images provided by the user. Our implementation requires only few parameters and is easy to handle by non-expert users via an intuitive graphical interface. It can be used for a multitude of applications and samples, provided the templates share sufficient pixel similarity with the objects in the images.

## Supplementary Information

See *Supplementary Material.pdf* and *Supplementary Figures.pdf* for further information. All data and software are archived on Zenodo (links in *Supplementary Material.pdf*).

## Acknowledgements

We thank Franz Schaefer (Children’s Hospital, Heidelberg) and Jochen Wittbrodt (COS, Heidelberg) for general support, Jakob Gierten (COS, Heidelberg) for sharing image data and Gunjan Pandey for proof-reading the manuscript. We are also grateful to the Fiji and KNIME communities for the support on their respective forum.

