## Supplementary Figures for "Multi-Template Matching: a versatile tool for object-localization in microscopy images"

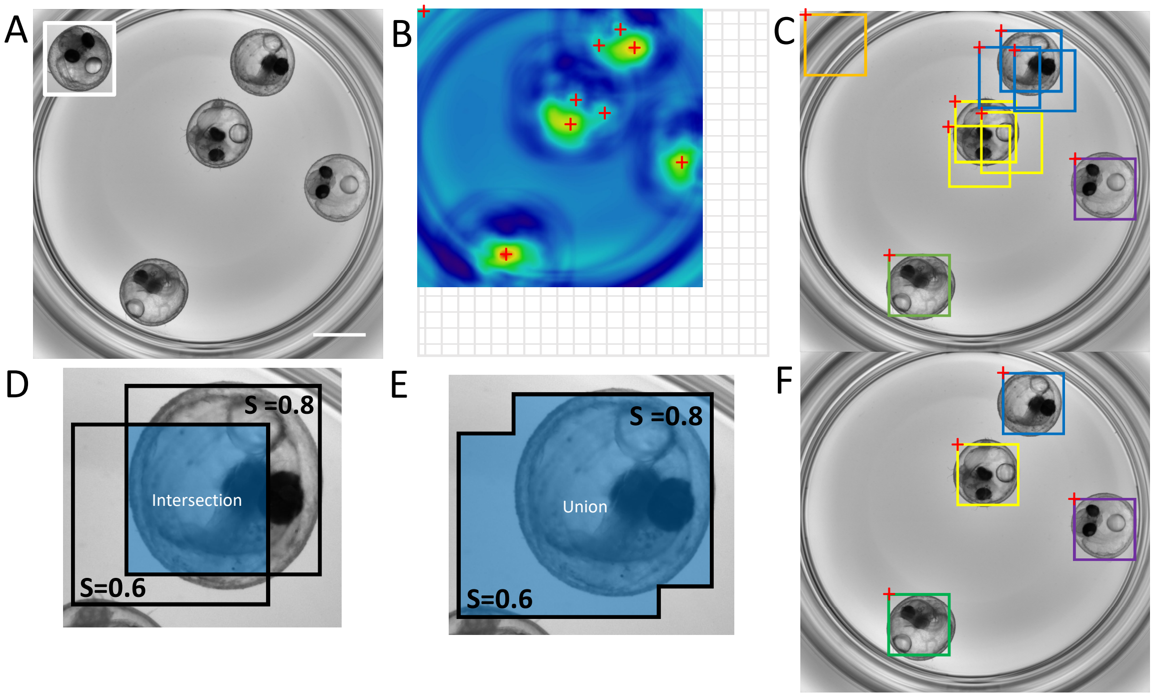


**Supplementary Figure 1: Multiple template matching on randomly oriented and positioned medaka embryos. (A)** Image in which the search is performed (2048x2048 pixels with scale bar of 1mm) and template as inset (400x414 pixels). The search was performed with a set of templates (original template, vertical and horizontal flip, each rotated by 90°, 180° and 270°). Parameters for the detection: score type: 0-mean normalized cross-correlation, score threshold: 0.35, Max overlap: 0.25. **(B)** One of the derived correlation maps from A: red crosses indicate possible local maxima before Non-Maxima Suppression (NMS). The grid area indicates the smaller size of the correlation map compared to the image in which the search is performed as explained in Supplementary material. **(C)** Bounding boxes associated to the maxima shown in B and overlaid on the searched image. Colours are highlighting overlapping bounding boxes. The bounding box dimensions are identical to the template used for the search. **(D,E)** Preventing overlapping detections by NMS. Shown are 2 overlapping bounding boxes predicting possible object locations. Each predicted location is associated to a probability score *S* to contain an object. The ratio between the **(D)** intersection and the **(E)** union area of the bounding boxes (Intersection over Union or IoU) is computed to decide whether the 2 overlapping bounding boxes are likely to predict the location of the same object (IoU close to 1) or the locations of distinct objects that are close to each other (IoU close to 0); for a detailed description of Non-Maxima Suppression see Supplementary Material. **(F)** Yielded object detections after NMS to return the N_objects=4 best detections.

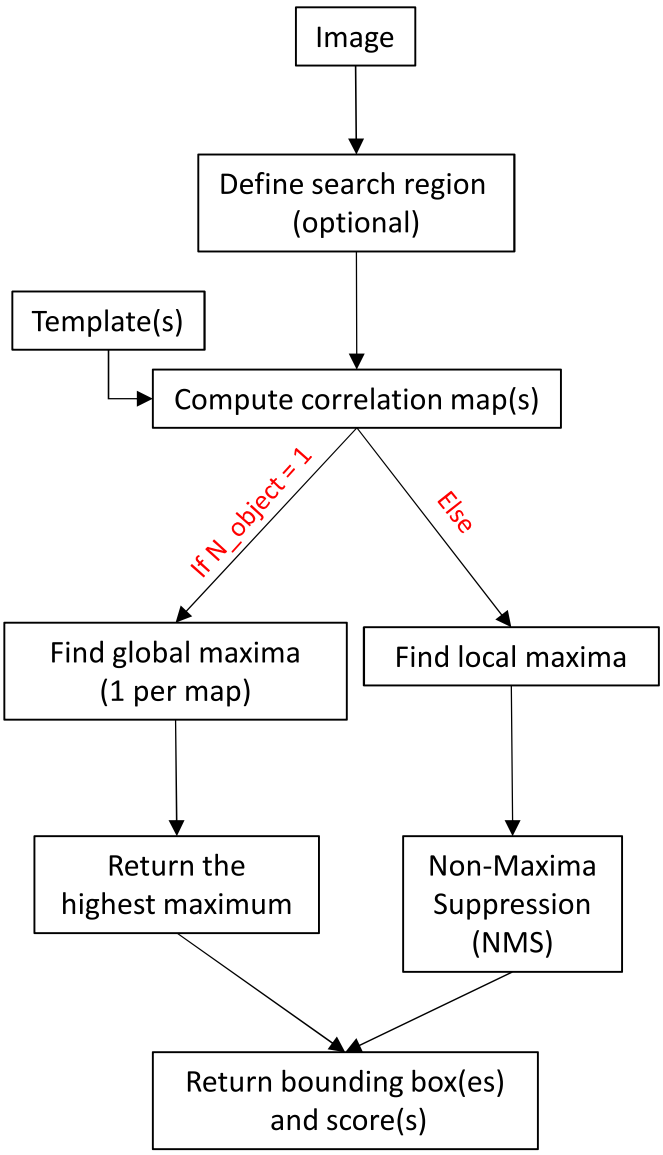


**Supplementary Figure 2: Flowchart of the implemented template matching**. The chart illustrates the sequential execution of the tool. For difference-based score, a difference map is computed, minima are detected instead of maxima and the lowest minima are returned. N_object is the expected number of objects in each image (provided by the user).

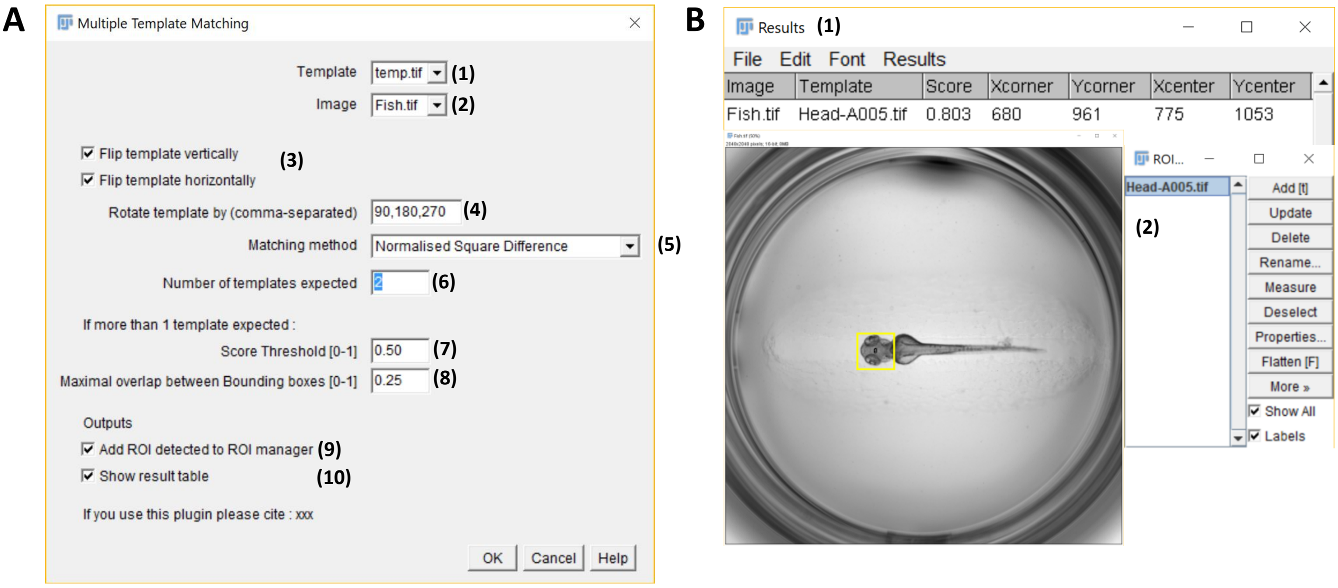


**Supplementary Figure 3: Implementation in Fiji. (A)** Graphical user interface with: (1) Dropdown menu to select the template image of the object of interest. The template must be smaller than the image specified in 2, (2) dropdown menu to select an image (or stack) in which to search for the template, (3) tick-boxes to optionally generate additional templates by horizontal/vertical flipping of the initial template, (4) input field for rotation angles to generate additional templates by rotations of the initial and, if selected, flipped templates. The angles are specified in degrees with clockwise orientation and must be separated by commas, (5) dropdown menu to choose the score used for the computation of the score map (normalised square-difference, normalised cross-correlation or 0-mean normalised cross-correlation), (6) input field to specify the number of templates/objects expected in the image, (7) input field to enter a score-threshold in the range 0-1. If the normalised square-difference is selected, only local minima with values below the threshold are returned. While for cross-correlation scores, maxima above this value are returned, (8) input field to specify the maximum value in range 0-1 for the intersection over union (IoU) between a pair of overlapping bounding boxes (Non-Maxima Suppression), (9) tick-box to select if the detected Regions Of Interest (ROI) should be added to Fiji ROI Manager, and (10) tick-box to specify if the result table containing coordinates and score of the predicted locations should be displayed at the end of the execution. Parameters 7 and 8 are only required if several objects are expected in each image. **(B)** Outputs of the plugin with (1) result table with each row showing the predicted object location containing the names of the image and template, the prediction score and coordinates of the top left corner and centre of the predicted bounding box, and (2) the detected ROI appended to the ROI Manager and highlighted on the image (yellow).

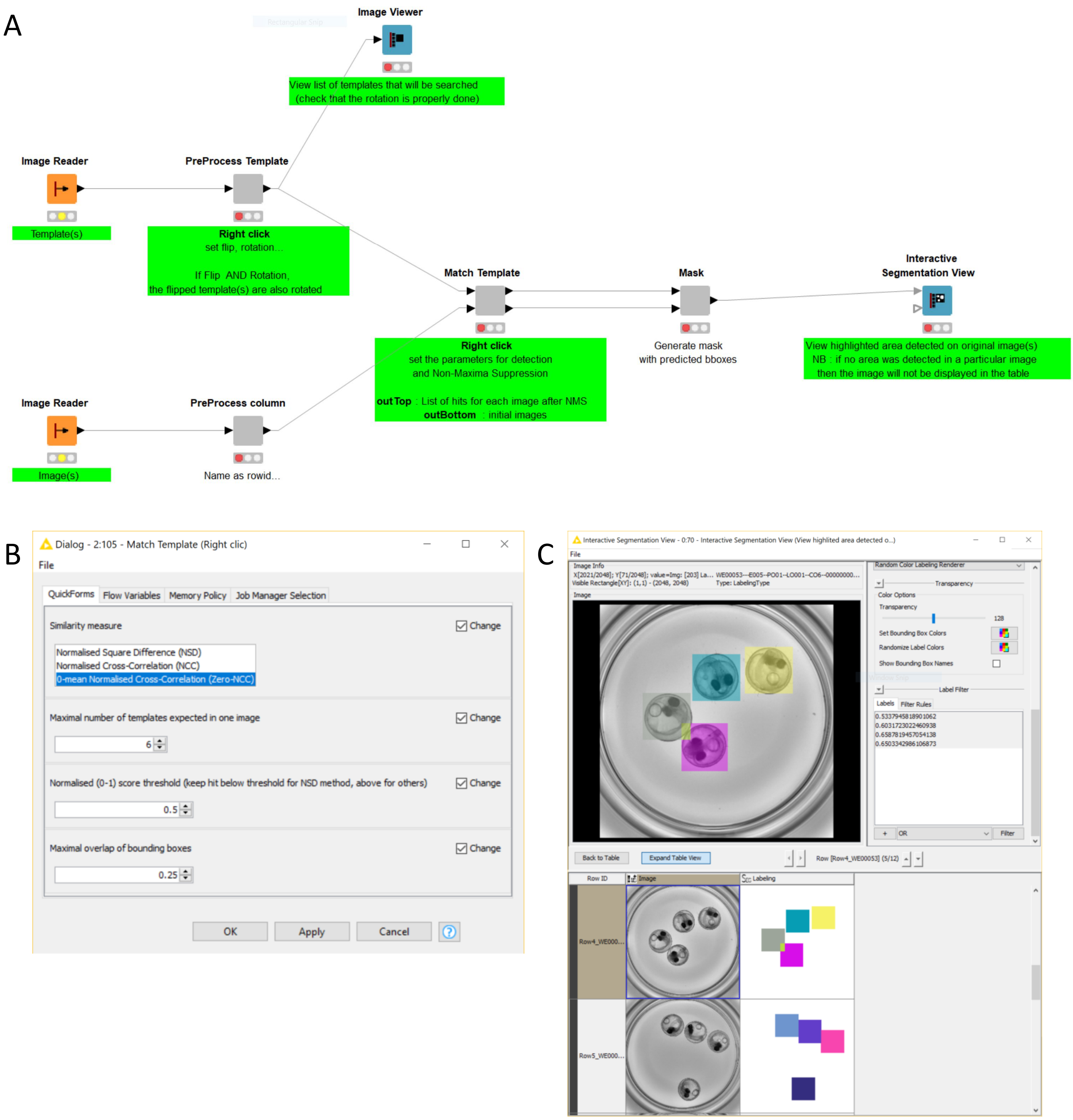


**Supplementary Figure 4: Implementation in KNIME. (A)** Screenshot of the KNIME workflow. The template and images are provided in the *Image Reader* nodes on the left side, the processing happens in the central metanode called '*Match Template*' containing a python node to interface with Python and OpenCV. The predicted locations can be visualised in the *Interactive Segmentation View* node on the right side (as shown in C). The user can interact with the nodes highlighted in green. **(B)** Graphical user interface of the central '*Match Template*' metanode for the configuration of the detection parameters, similarly to the Fiji implementation (see Suppl. Fig. 3A). **(C)** Predicted locations as viewed in the *Interactive Segmentation View* node. The predicted locations form a mask image that can be visualised as overlay on the image. A result table containing the bounding box location is also provided (not shown).

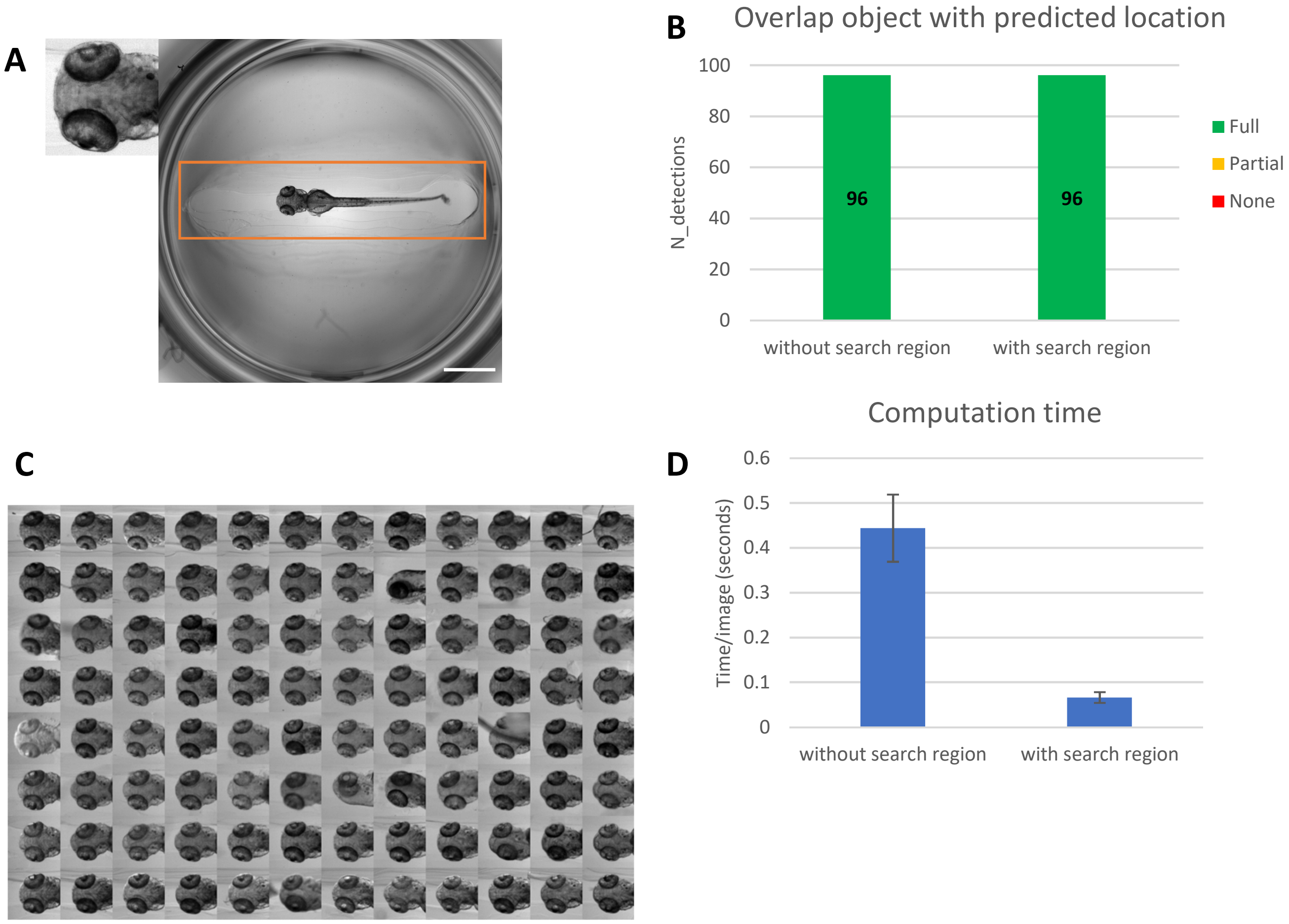


**Supplementary Figure 5: Template matching for head region detection in oriented zebrafish larvae. (A)** Single template (188x194 pixels, no additional transformation) and image (2048x2048 pixels, scale bar of 1mm) in which the search is performed. The orange rectangle shows the optionally used restricted search region (1820x452). Parameters for the detection: score type: 0-mean normalised cross-correlation with 1 expected object/image. **(B)** Result of the detection for N=96 images, with and without search region (both 100% detection rate). *Included/Partial/Missed* in figure legend refer to the position of the object within the predicted bounding box. **(C)** Montage of detected zebrafish larval head regions within a 96 well plate (as in Fig. 1C). **(D)** Mean computation time per image (error bars show standard deviation) for the different conditions as in B using the same computing hardware as in the main text. Prior information about the position of the sample within the field of view (e.g. due to standardized sample mounting) can be used to specify a search region, drastically accelerating the computation and reducing the chance of incorrect predictions.

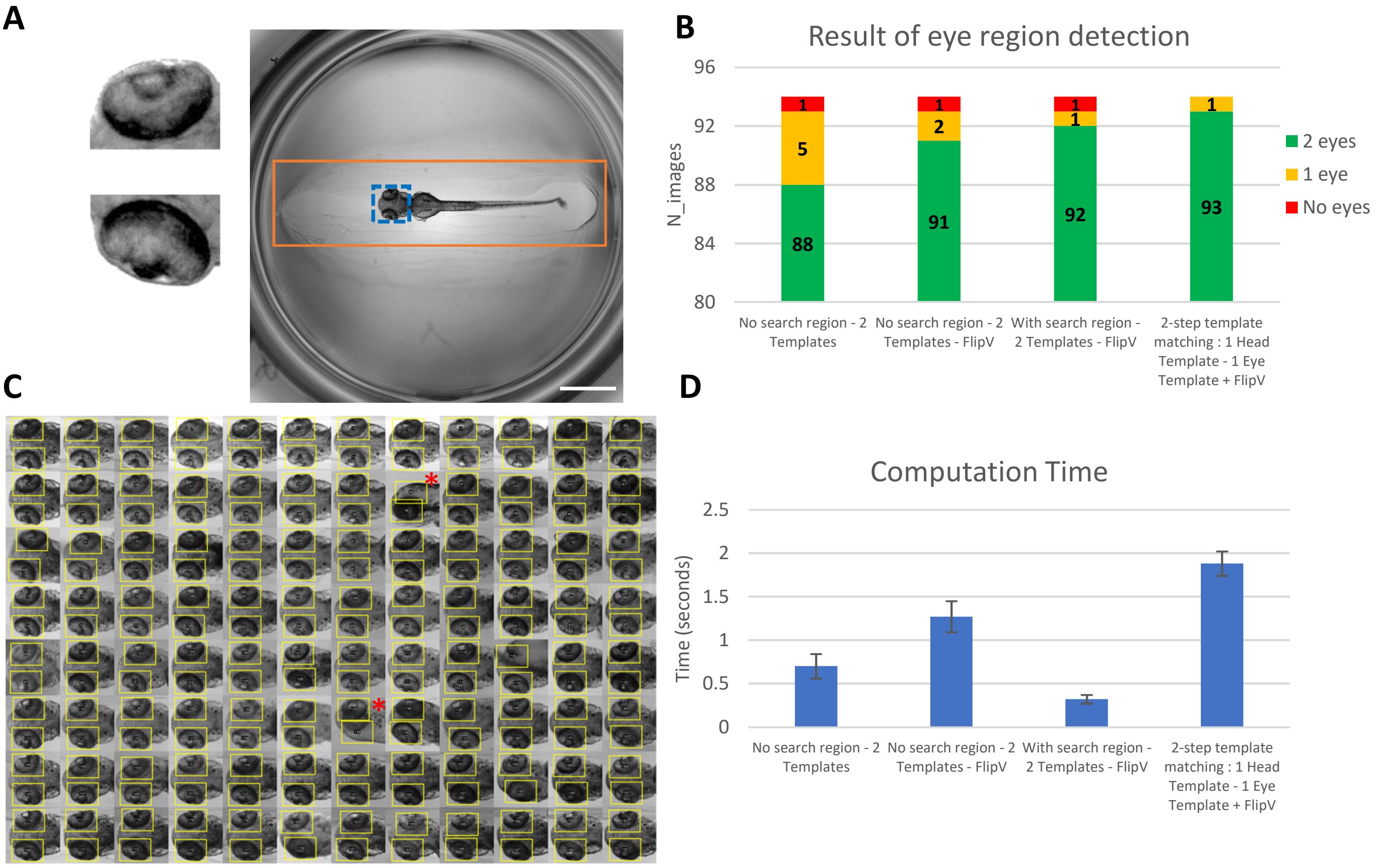


**Supplementary Figure 6: Template matching for eye region detection in oriented zebrafish larvae. (A)** Templates (108x76 pixels) and image in which the search is performed (2048x2048 pixels, scale bar of 1 mm). Orange rectangle indicates optional search region and blue dotted rectangle the head region template used for 2-step template matching (see B and D). Parameters for the detection: Vertical flipping of the templates (FlipV), score type: 0-mean normalized cross-correlation, score threshold: 0.5, max overlap: 0.25 **(B)** Result of the detections for N=94 images. *2 eyes/1 eye/no eye* in figure legend refer to the outcome of eye region detection in each larvae. Vertical flipping of the templates readily increases the number of genuine matches. The 2-step template matching approach (search of template within a previously identified ROI) offers the best results and is recommended for more challenging template images (see Supplementary Macro). **(C)** Montage of the eye regions detections (yellow) for the 2-step matching approach as in B and D. Specimen marked with a red asterisk are excluded from the count in B as they are not dorsally oriented. **(D)** Mean computation time per image (error bars show standard deviation) for the different conditions (as in B) using the same hardware as in the main text.

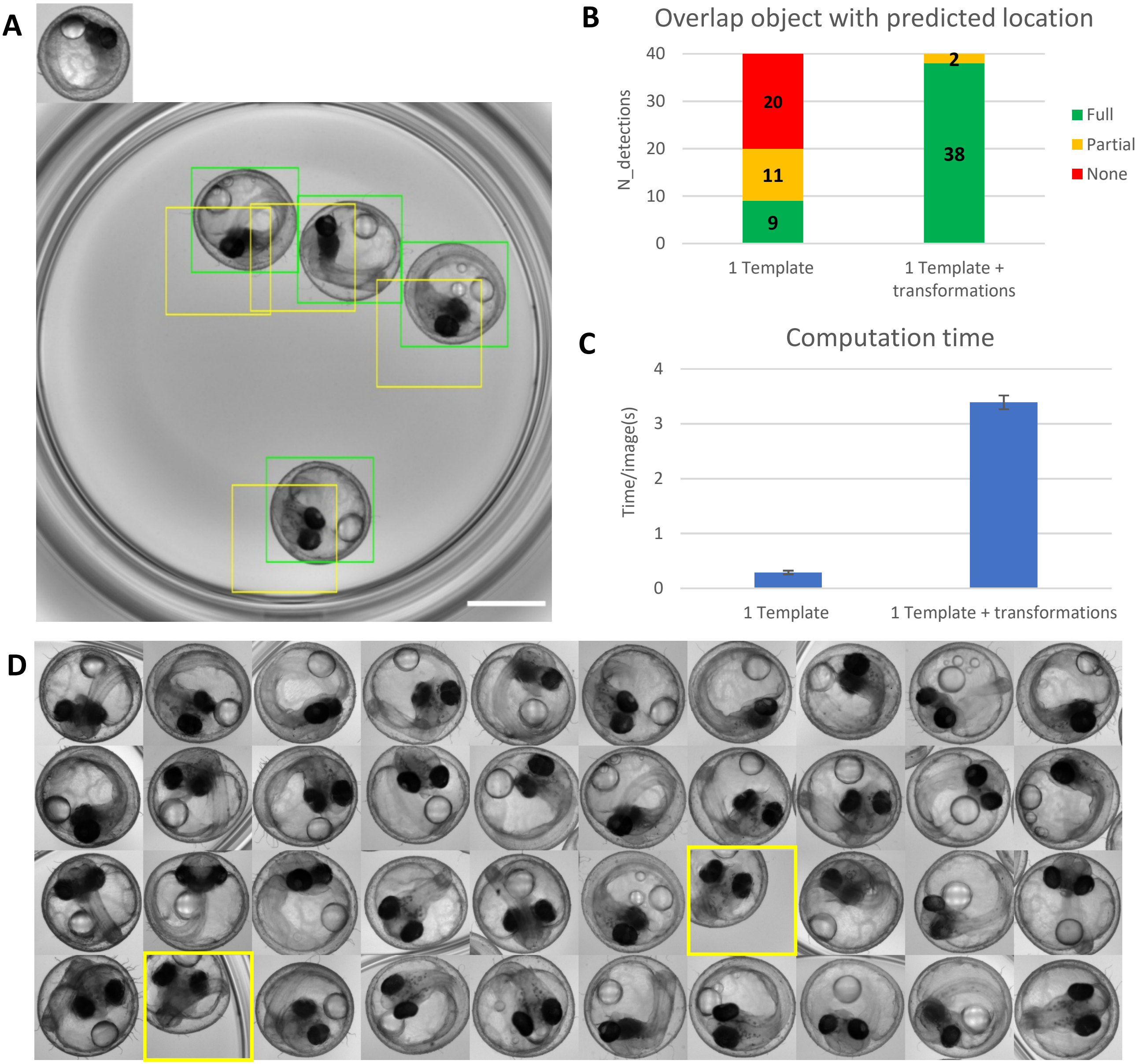


**Supplementary Figure 7: Template matching for the localization of randomly oriented and positioned medaka embryos.** **(A)** Initial template (410x420 pixels) and one of the image in which the search is performed (2048x2048 pixels, scale bar of 1 mm). The yellow bounding boxes indicate predicted locations when only the original template in A is used, the green boxes indicate predicted locations when using a set of templates (original, horizontal and vertical flipping, rotation of the original and flipped templates by 90°,180° and 270°). Parameters for the detection: score type: 0-mean normalized cross-correlation, score threshold:0.35, Max overlap:0.25. **(B)** Result of the detections for 10 images each containing 4 embryos (i.e. 40 embryos in total). *Included/Partial/Missed* in figure legend refer to the position of the object within the predicted bounding box. **(C)** Mean computation time per image (error bars show standard deviation) for the different conditions using the same hardware as in the main text. The computation time for each image scales linearly with the number of templates. **(D)** Montage of the detected regions for 10 images similar to A each containing 4 embryos (1 column/image). The montage corresponds to the benchmark *"1 Template+transformations"* as in B and C. Yellow bounding boxes indicate the 2 detections classified as *Partial.*
