## Supplementary Material for "Multi-Template Matching: a versatile tool for object-localization in microscopy images"

[1] Acquifer is a division of Ditabis, Digital Biomedical Imaging Systems AG, Pforzheim, Germany.

[2] Centre of Pediatrics and Adolescent Medicine, University Hospital Heidelberg, Germany

### Link to online supplementary resources

**- GitHub repository for the Fiji implementation (including supplementary macro for 2-step template matching): https://github.com/LauLauThom/MultiTemplateMatching**

**- GitHub repository for the KNIME implementation: https://github.com/LauLauThom/MultipleTemplateMatching-KNIME**

**- KNIME workflow available for download on** nodepit space and on the GitHub repository above

**- Dataset used for Fig.1, Suppl.Fig.1 and 7**

Gierten Jakob, Gehrig Jochen, & Thomas Laurent. (2019). 102 hpf medaka embryos in 96 well plate (4 embryo/well) - brightfield - 2X magnification - ACQUIFER Imaging Machine (Version 1) [Data set]. Zenodo. **http://doi.org/10.5281/zenodo.2650147**

**- Dataset used for Fig.1, Suppl.Fig.5 and 6**

Gehrig, Jochen. (2019). 3dpf zebrafish larvae, 96 well plate,Tg(wt1b:EGFP), dorsal view, ACQUIFER Imaging Machine [Data set]. Zenodo. **http://doi.org/10.5281/zenodo.2650162**

**- Video tutorials for Fiji and KNIME implementations**

**On Zenodo:** Thomas Laurent. (2019). Video tutorial for Multi-Template-Matching implementation in Fiji and KNIME (Version 1). Zenodo. **http:// doi.org/10.5281/zenodo.2650171**

**On Youtube:**

1) Installation in Fiji and single object detection: **https://youtu.be/KlzIqSG5XBU**
2) Multiple objects detection (Fiji): **https://youtu.be/-PoZihjJIjQ**
3) Macro recording (Fiji): **https://youtu.be/tTonuVgk2e0**
4) KNIME implementation **https://youtu.be/pldrWMJhE3o**

**- Pre-configured Fiji installation**

Thomas Laurent, & Gehrig Jochen. (2019, April 25). Archived preconfigured Fiji installation for multiple template matching (Version 1.0). Zenodo. **http://doi.org/10.5281/zenodo.2650856**

**- Pre-configured KNIME installation**

Laurent Thomas. (2019, April 25). Archived KNIME and anaconda python environment for multiple-template-matching (Version 1). Zenodo. **http://doi.org/10.5281/zenodo.2650851**

### Requirements

At the time of the publication, the Fiji plugins were tested with Fiji running ImageJ 1.52k, Java 1.8.0_66 and IJ-OpenCV 1.2.1.

The KNIME workflow was tested with KNIME 3.7.1, Python 3.6.8 and OpenCV 3.4.2.

### Detailed description

#### Template Matching

Template matching is an iterative search algorithm that uses the template image as a rectangular search window sliding over the target image. For every *(x, y)* position of this sliding window in the target image, the algorithm computes a similarity score as the sum of the pixel intensity differences or correlations between the overlapping template pixels and current image-patch pixels. This similarity score is assigned to the pixel at position *(x, y)* in the resulting score map. This procedure is repeated for every *(x, y)* position of the sliding window in the image, that offers full overlap between the template and an underlying image-patch. Therefore, the resulting score map between a template of dimensions ($W_{Template}$, $H_{Template}$) and an image of dimensions ($W_{Image}$, $H_{Image}$) is an image of dimensions ($W_{Image}$-$W_{Template}$, $H_{Image}$-$H_{Template}$).

In the case of a difference-based score, a high probability to find the template *T* at a position *(x, y)* in the image *I* corresponds to a low grey value for the pixel at position *(x, y)* in the score map, while for the correlation-based scores it corresponds to a high grey value for that pixel. Therefore, the possible positions of the template in the image are provided by the ($x_M$, $y_M$) coordinates of the extrema in the score map (minima or maxima depending on the score), while the pixel value of each extrema corresponds to the associated score.

If a single object is expected in the image, a global extrema detection is performed on the score map to get the predicted position of the template. If several objects are expected, local extrema with a score above a user-defined threshold are collected and subsequently filtered by Non-Maxima Suppression (NMS, see section below) to remove predictions in too close vicinity. The NMS is terminated once the N highest score predictions have been collected (with N the expected number of objects in the image) or when there are no further extrema to test for NMS. In this latter case, the algorithm will return less than N predicted locations for the template image.

The resulting predictions can be visualized as a set of bounding boxes overlapped on the original image, corresponding to the expected template locations. The bounding boxes have identical dimensions to the template used for the search and are positioned by placing the top-left pixel to position ($x_M$, $y_M$) in the image.

Similarly, an image can be searched using several templates (e.g. additional geometrical transformations of the template, phenotypic variations a biological object, several biological objects etc.) to maximize the probability of detecting objects. Each sequential search with a different template yields a new score map and a set of extrema. The global extrema detection or NMS is in this case performed on the union of the different extrema sets.

**Choice of the score**

Normalized scores are used for the computation of the score map. This has a number of advantages: (i) it prevents the score to be biased by very bright or dark pixel values of either the template or the image, (ii) it yields a score map with pixel values ranging from 0 to 1, which facilitates the application of a threshold on the score for the local extrema detection, (iii) it also ensures that score maps generated from different templates are still comparable.

The *matchTemplate* function from OpenCV is available with different types of scores (see **OpenCv documentation**).

For the normalization of the score, the sum of the pixel value differences (for difference-score) - or products (for correlation score) - between the template and the image patch are divided by a term proportional to the pixel intensities of both the template and the current image-patch.

With the *0-mean normalized cross-correlation*, the mean grey value of the template and of the image-patch are subtracted from each of their pixel respectively before the computation of the normalized-correlation. This has an additional intensity normalization effect which makes the detection particularly robust to local illumination changes in the image as it can often occur in microscopy. This choice of score thus usually yields the best predicted locations and is proposed as the default choice.

**Non-Maxima Suppression**

As discussed above, the possible template locations in an image can be deducted from the positions of the extrema in the score map. When only one template object is expected in the image, the position of the global extremum directly yields the expected template image location. However, when several objects should be detected in the image, one needs to first identify the locations of the local extrema. These are identified as the extremum of a small neighborhood (like a 3x3 pixel window centered on the extremum) with a score value above (or below) a threshold. In some implementations, the size of the neighborhood can be adjusted to avoid too close local extrema that would correspond in our case to overlapping detections of the same object, as it usually occurs with object-recognition algorithms. As the *(x, y)* coordinates of an extrema indicate the position of the top left pixel of the bounding box predicting the object location, it is virtually impossible to define a neighborhood-size around that corner that would prevent overlapping detections while conserving genuine detections of objects close to each other. Therefore, we use another strategy for Non-Maxima Suppression based on the degree of overlap between neighboring bounding boxes similar to (Alexe *et al.*, 2012) (Supplementary Figure 1.D-E). It is based on the computation of the ratio between the Intersection area and Union area (referred to as Intersection over Union or IoU) for a pair of bounding boxes. Basically, if the bounding boxes highly overlap, the ratio will be close to 1, while if the overlap is small the ratio will tend to 0. In practice, a user-defined threshold on the IoU is used as the maximal value of overlap allowed between bounding boxes, such that a bounding box overlapping above the IoU threshold with another bounding box of higher score is discarded. This effectively removes overlapping detection of the same object, while still allowing the detections of close objects for which the bounding boxes might slightly overlap.

The IoU has the advantage to be comparable for any pair of bounding boxes, independently of their dimensions as it is normalized by the union of the area.

### Implementation

We implemented our multiple-template matching pipeline both in Fiji and KNIME using the jython and python languages respectively. The computation of the score map given a template and a test image is provided by the function *matchTemplate* from the OpenCV library. This function is defined for greyscale 8-bit or 32-bit images only, therefore, to be compatible with images of any bit depth, the 16-bit and RGB images are automatically converted to 32-bit internally in our implementation.

OpenCV is distributed as a C++, Python and Java library. In Fiji, functions from the OpenCV package can be easily accessed by enabling the IJ-OpenCV update site (Domínguez et al., 2017).

Our Fiji macro can similarly be installed by enabling the Multi-Template-Matching update site, while for KNIME we provide the workflow files and a dependency file to setup the associated python environment. All source codes can be found on the GitHub repositories: *LauLauThom/MultiTemplateMatching* and *LauLauThom/MultipleTemplateMatching-KNIME*.

**Template pre-processing**

The workflow/plugin can take as input argument a list of templates and a list of images for which to search for the template(s). The template(s) can additionally be flipped (vertically and/or horizontally) and/or rotated by a user-provided list of angles within the same interface. If flipping and rotation are selected, each template in the list will be flipped and both the original and flipped versions of the template will be rotated by the same angles. For a given rotation angle, an image and its flipped version will not result in the same image except in case of symmetry along the flipping axis.

Therefore, for an initial list of $T$ templates, if both vertical and horizontal flipping are selected and k additional rotations are performed, we end up with a final list of $T' = T x 3 x (k+1)$ templates.

Since the template matching algorithm can only search for rectangular templates, the templates rotated by non-quadrants rotations results in larger rectangular images with some empty areas. To prevent those areas from penalizing the score, they are filled with the modal grey value of the original template for the Fiji implementation or with the value of the pixel at the border of the original template for the KNIME implementation. It is important to note that the portion of empty areas is larger when rotating templates of high aspect ratio.

**Extrema Detection**

If the expected number of objects per image $N$=1, a global extremum detection (minimum for difference-based score, maximum for correlation-based score) is performed in each of the $T'$ score maps resulting in a set *S* of *T'* extrema. The predicted location of the template is then given by the extrema of highest or lowest score in *S* (depending on the choice of score: correlation/difference).

If *N>1*, a local extrema detection is performed on each score map with a threshold on the score. For correlation-based scores, local maxima with a score above the threshold are returned while for the difference-based score, the local minima are detected similarly as the local maxima of the inverted difference map (1-Difference Map). The local extrema from the different score maps are then merged into a large set *S*. NMS is finally performed on *S* to yield up to *N* best non-overlapping detections.

**Non-Maxima Suppression (NMS)**

We wrote our own implementation of NMS, considering a threshold on the IoU but also the expected number of objects per image *N*. Our NMS will thus return from a pool of possible template locations, up to *N* best locations for which the predicted bounding boxes do not overlap above the specified IoU threshold.

As discussed above, each extremum of a score map is associated to a bounding box of dimensions identical to the template used for the computation of that score map, and the associated score is the pixel value of the extrema in the score map image.

For correlation-based score the detailed procedure for the NMS is as following (for difference-based score the procedure is the same but with the opposite sorting of scores). We first order the maxima in *S* by descending scores (highest scores first). We then initialize the final set of maxima S' with the highest maximum in *S*. After that we iterate over the remaining maxima in *S*, and for each of them compute the IoU between its associated bounding box and the bounding boxes of the previously collected hits in S'.

If the list of IoU are all below the threshold, which means that the current maxima is not substantially overlapping with any maxima of higher score in *S'*, then the tested maxima is added to *S'* otherwise the iteration continues with the next maxima in *S*, until we have collected *N* maxima in *S'* or until there are no more maxima available to test in *S*.

### Example applications and benchmarking

We tested our implementation for the detection of single or multiple objects, using either one template image (Suppl.Fig.5), distinct template images (Suppl.Fig.6) or one template image and a set of transformed versions of it (Suppl.Fig.7).

We also show the result for a custom-made 2-step template matching using the ImageJ macro language in Supplementary Figure 6. The principle of the 2-step template matching is to perform a first template matching to robustly detect a region, that is then successively used as limiting search region for a second template matching with a smaller template. This approach can yield better detections than the 1-step version, especially for the detections of small regions with homogeneous texture.

We report for each benchmark the localization results and the average computation time per image in seconds.

All benchmarks were carried out with the plugin named “Template Matching Folder” using the same hardware configuration as mentioned in the main text.
